# The SUMO Protease SENP3 regulates Mitochondrial Autophagy mediated by Fis1

**DOI:** 10.1101/685834

**Authors:** Emily Waters, Kevin A. Wilkinson, Ruth E. Carmichael, Chun Guo

**Affiliations:** Department of Biomedical Science, University of Sheffield, Firth Court, Western Bank, Sheffield, S10 2TN, U.K; School of Biochemistry, Medical Sciences Building, University of Bristol, University Walk, Bristol, BS8 1TD, U.K

**Keywords:** SENP3, Fis1, SUMO, Organellar Stress, Mitophagy

## Abstract

Mitochondria are unavoidably subject to organellar stress resulting from exposure to a range of reactive molecular species. Consequently, cells operate a poorly understood quality control programme of mitophagy to facilitate elimination of dysfunctional mitochondria. Here we use a model stressor, deferiprone (DFP), to investigate the molecular basis for stress-induced mitophagy. We show that mitochondrial fission 1 protein (Fis1) is required for DFP-induced mitophagy and that Fis1 is SUMOylated at K149, an amino acid residue critical for Fis1 mitochondrial localization. We find that DFP treatment leads to the stabilisation of the SUMO protease SENP3, which is mediated by downregulation of the E3 ubiquitin (Ub) ligase CHIP. SENP3 is responsible for Fis1 deSUMOylation and depletion of SENP3 abolishes DFP-induced mitochondrial mitophagy. Furthermore, preventing Fis1 SUMOylation by conservative K149R mutation enhances Fis1 mitochondrial localization. Critically, expressing a Fis1 K149R mutant restores DFP-induced mitophagy in SENP3 depleted cells. Thus, we propose a model in which SENP3-mediated deSUMOylation facilitates Fis1 mitochondrial localization to underpin stress-induced mitophagy.

## Introduction

Accurate and proper degradation of dysfunctional mitochondria by mitophagy is essential for maintaining control over mitochondrial quality and quantity. However, mitophagic mechanisms, and their control, remain unclear (Green, Galluzzi et al., 2011). Degradation of dysfunctional mitochondria occurs via a poorly understood quality control program involving mitochondrial fission of dysfunctional components (Twig, Elorza et al., 2008), their accumulation in autophagosomes (Jimenez-Orgaz, Kvainickas et al., 2018), and subsequent lysosomal fusion resulting in degradation (Sun, Yun et al., 2015).

Accumulation of dysfunctional mitochondria is associated with both aging (Biala, Dhingra et al., 2015, Gonzalez-Freire, de Cabo et al., 2015, Green et al., 2011, Lopez-Otin, Blasco et al., 2013) and age-related diseases (Lionaki, Markaki et al., 2015, Quiros, Langer et al., 2015, Wong & Holzbaur, 2015). Mitochondrial fission ensures efficient mitophagy (Archer, 2013; Burman et al., 2017; Mao et al., 2013; Twig et al., 2008) and impairment of mitochondrial fission is evident in aging (Morozov et al., 2017) and age-related diseases (Chen and Chan, 2009; Lionaki et al., 2015; Zhang et al., 2016). Mitochondrial fission mainly depends on the GTPase Drp1 (Frank et al., 2001), which is recruited to the mitochondrial surface and acts by engaging with specific mitochondrial docking proteins such as Mff, MID49, MID51 and probably Fis1 in mammalian cells (Loson et al., 2013). However, Drp1 itself does not appear to be essential for mitophagy (Mendl, Occhipinti et al., 2011, Song, Gong et al., 2015, Yamashita, Jin et al., 2016). More recently, evidence has emerged for a major role of Fis1 in the disposal of defective mitochondria induced by stressors, including antimycin A, paraquat and deferiprone (DFP), which regulates the formation of reactive oxygen species (Huang and Chan, 2016; Rojansky et al., 2016; Shen et al., 2014; Yamano et al., 2014). Fis1 has been identified, though not validated, in a proteomics screen as a target for post-translational modification (PTM) by Small Ubiquitin-like MOdifier protein (SUMO) *in vivo* (Tirard et al., 2012).

SUMOylation involves the reversible covalent conjugation of SUMO to specific lysine(s) in target proteins (Boddy, Howe et al., 1996, Kamitani, Nguyen et al., 1997, Shen, Pardington-Purtymun et al., 1996). Three distinct *SUMO* gene products in mammals are validated for conjugation: SUMO-1 shares ∼50% sequence homology with SUMO-2 and −3, which differ by only 3 amino acids and are collectively referred to as SUMO-2/3. SUMO conjugation is highly transient and is reversed by the action of SUMO specific proteases (Henley, Craig et al., 2014, Wilkinson & Henley, 2010). SUMO proteases deconjugate SUMO from SUMOylated proteins in a process termed deSUMOylation (Hickey, Wilson et al., 2012, Nayak & Muller, 2014). The extent of target protein SUMOylation in any given state arises from the balance of the opposing activities of SUMO conjugating and deconjugating enzymes. To date, three families of SUMO proteases have been identified: deSUMOylating isopeptidase 1 and 2 (DeSI1 and DeSI2), Ubiquitin-specific protease-like 1 (USPL1), and Sentrin-specific proteases (SENPs). The largest and best characterised family of SUMO proteases, SENPs consist of six cysteine proteases (SENP1-3 and 5-7), each having a distinct subcellular localisation and SUMO isoform substrate preference (Guo & Henley, 2014, Hickey et al., 2012).

SENP3 is an essential deSUMOylating enzyme primarily responsible for deconjugating SUMO-2/3 from modified proteins, as evidenced by the findings that SENP3 depletion results in a significant increase in global SUMO-2/3-ylation in mammalian cells (Fanis, Gillemans et al., 2012, Guo, Hildick et al., 2013, Haindl, Harasim et al., 2008, Luo, Gurung et al., 2017). Although SENP3 is known to play key roles in many processes such as inflammation (Huang, Ghisletti et al., 2011), stem cell differentiation (Nayak, Viale-Bouroncle et al., 2014), ribosome biogenesis (Raman, Weir et al., 2016) and cell stress responses (Guo et al., 2013, Huang, Han et al., 2009, Yan, Sun et al., 2010), specific targets and physiological roles for this enzyme are largely unknown. Previously believed to be an exclusively nuclear protein, our recent work has revealed that, importantly, SENP3 also resides in the cytoplasm, and is required for mitochondrial fission (Guo et al., 2013, Guo, Wilkinson et al., 2017), a process which ensures mitophagy-mediated removal of damaged or dysfunctional mitochondria (Twig et al., 2008). Moreover, emerging evidence has strongly implicated SENP3 in ageing and age-related degenerative diseases. For example, *SENP3* gene expression is markedly reduced in brain samples either from healthy aged subjects (Lu, Pan et al., 2004) or from Alzheimer’s disease patients (Weeraratna, Kalehua et al., 2007), and SENP3 levels are altered in brain samples from the patients of Down’s syndrome (Binda, Heimann et al., 2017) which is associated with premature aging and is a risk factor for dementia development (Roth, Sun et al., 1996, Weksler, Szabo et al., 2013). However, the functional consequences and molecular mechanism(s) underlying the changes in SENP3 levels remain undetermined. Furthermore, accumulation of dysfunctional mitochondria is linked with aging and age-related degenerative diseases, and this is believed to be due to defects in a mitochondrial quality control program enforced by mitophagy, suggesting the possible involvement of SENP3 in mitochondrial quality control.

Here, we have investigated the role of SENP3 in mitophagy-mediated mitochondrial disposal in model (HeLa) cells by investigating the effects of altered SENP3 levels on induced mitophagy and autophagosome formation. Cells were exposed to a model stressor, the iron chelator DFP, which specifically induces mitophagy, but not general autophagy (Allen, Toth et al., 2013, Sargsyan, Cai et al., 2015, Yamashita et al., 2016). DFP is used for treatment of iron overload in inherited and degenerative diseases such as Thalassemia major and Parkinson’s disease. We show that levels of the SUMO-2/3-specific deSUMOylating enzyme SENP3 are greatly stabilised upon DFP treatment via a pathway that involves the reduction in levels of the E3 ubiquitin (Ub)-protein ligase CHIP. This stabilisation of SENP3 decreases Fis1 SUMO-2/3-ylation, enhances Fis1 mitochondrial localization and induces Fis1-mediated autophagosome formation and mitochondrial disposal. Taken together, our results reveal a novel mechanism dependent on the presence of SENP3 in co-ordinated regulation of the SUMOylation status of Fis1 to ensure mitochondrial quality control.

## Results

### DFP-induced mitophagy is accompanied by decreased SUMO-2/3 conjugation, increased SENP3 levels and reduced CHIP levels

DFP is known to induce Parkin-independent mitophagy (Allen et al., 2013). Consistent with this, we detected induction of the autophagosomal marker LC3-II by DFP in HeLa cells that express little or no Parkin (Supplementary Figure 1A). To detect lysosomal-mediated acidification of engulfed mitochondria, we designed and characterised a novel probe termed MitoPHfluorin (Supplementary Figure 1B). This probe consists of mCherry–SEP (a pH-sensitive GFP-variant (Ashby, Ibaraki et al., 2004) tagged to the mitochondrial targeting sequence of the ActA protein of *Listeria monocytogenes* (Pistor, Chakraborty et al., 1994), resulting in a quenchable dual-fluorescence mitochondrial marker (Supplementary Figure 1B). HeLa cells treated with DFP show increased levels of SEP fluorescence quenching that results in the appearance of red puncta (Supplementary Figure 1C), indicative of elevated autolysosome formation.

Interestingly, DFP treatment also led to a significant decrease in global SUMO-2/3-ylation (Figure 1A) but not SUMO-1-lyation (Figure 1B) in HeLa cells. Moreover, levels of the deSUMOylating enzyme SENP3 (Figure 1C), but not SENP5 (Figure 1D), were increased by DFP. Interestingly DFP treatment decreases the levels of CHIP (Figure 1E), an E3 Ub ligase known to control SENP3 levels under basal conditions (Yan et al., 2010). Consistent with previous findings, knocking down CHIP in HeLa cells increased the levels of SENP3 but not SENP5 (Figure 1F), and prevented the DFP-induced increase in SENP3 levels (Figure 1G). Taken together, these findings suggest that the downregulation of CHIP upon DFP treatment underlies increased SENP3 levels, thereby modulating levels of cellular SUMO-2/3 conjugation.

**Figure 1.**
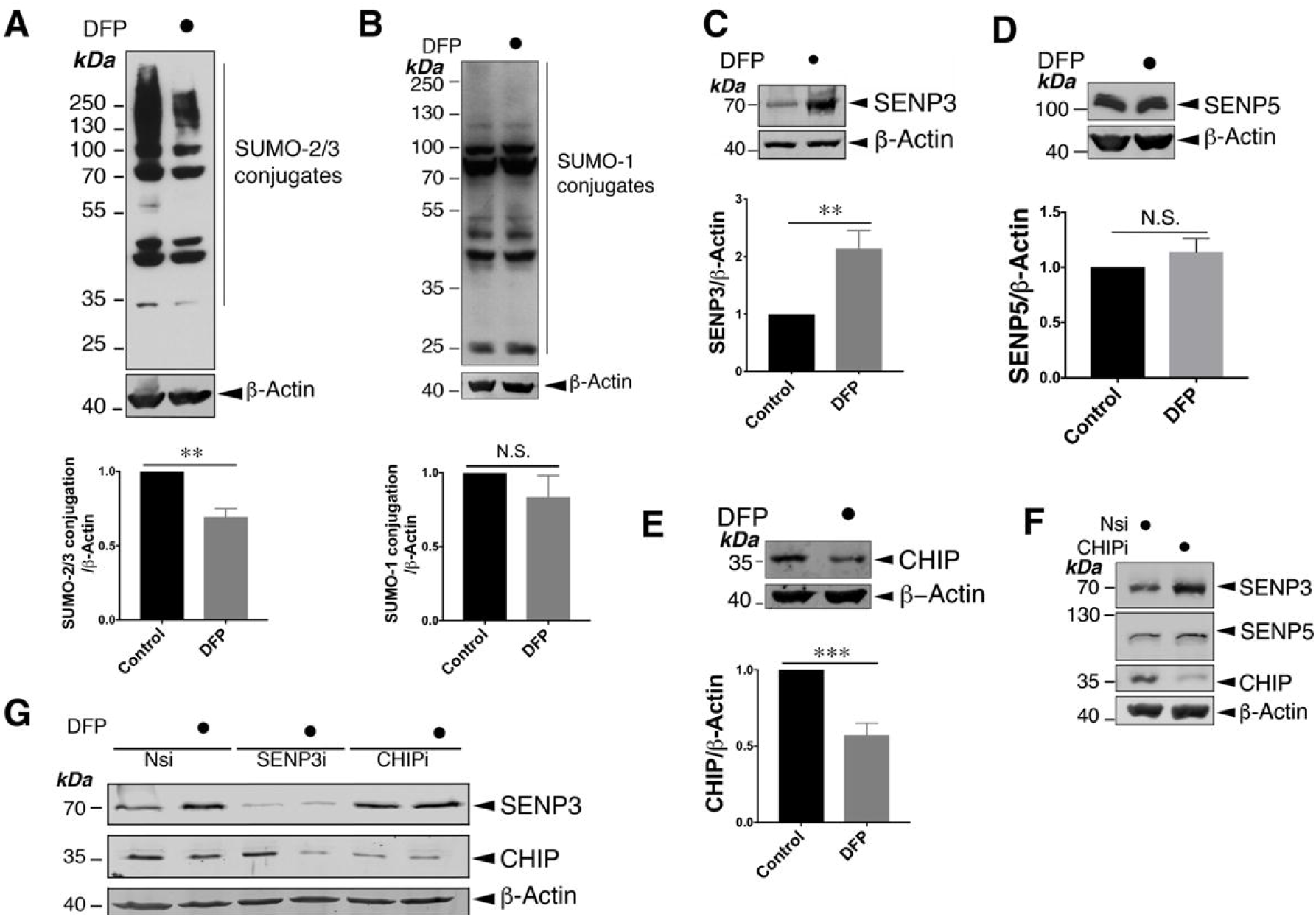
Iron chelation induces protein deSUMO-2/3-ylation coinciding with SENP3 stabilisation and CHIP reduction. A. Treatment of HeLa cells with DFP (1mM for 24 h) reduces the conjugation of SUMO-2/3 (n=5; **, p<0.01; paired t-test). B. Treatment of HeLa cells with DFP (1mM for 24 h) does not change global SUMO-1-ylation n=5; p>0.05; N.S., not significant; paired-t test). C. Treatment of HeLa cells with DFP (1mM for 24 h) leads to increased levels of SENP3 (n=8, **p<0.01; paired t-test). D. Treatment of HeLa cells with DFP (1mM for 24 h) does not change levels of SENP5 (n=5; p>0.05; N.S., not significant; paired-t test). E. Treatment of HeLa cells with DFP (1mM for 24 h) decreases levels of CHIP (n=8, ***p<0.001; paired t-test). In (A)∼(E) values are presented as mean ± SEM and are normalised to the control value. F. CHIP knockdown increases levels of SENP3 but not SENP5 in HeLa cells. Nsi or CHIPi (Nsi, non-specific siRNA; CHIPi, CHIP siRNA; concentration, 20nM) was transfected into HeLa cells for 48 h. Lysate samples were blotted as indicated. G. CHIP knockdown prevents the DFP-induced increase in SENP3 levels. Nsi, SENP3i or CHIPi (Nsi, non-specific siRNA; SENP3i, SENP3 siRNA; concentration 20nM; CHIPi, CHIP siRNA; concentration, 20nM) were transfected into HeLa cells for 48 h. Two days post transfection and the cells were treated with DFP for a further 24 h. In (A)∼(G) whole cell lysate samples were blotted as indicated.

### SENP3 is required for mitophagy induced by DFP

Mitophagy-mediated disposal of dysfunctional mitochondria occurs via a multi-step process involving two intracellular events: mitochondrial autophagosome formation and subsequent autolysosome formation (Yamashita & Kanki, 2017). To explore if SENP3 is involved in DFP-induced mitochondrial autophagosome formation, we examined the effect of RNAi-mediated knockdown of SENP3 on the expression of the autophagosomal marker LC3-II in HeLa cells treated with DFP. SENP3 knockdown significantly reduced DFP-induced LC3-II expression (Figure 2A), indicating an essential role for SENP3 in autophagosome formation. We then compared mitochondria-containing autolysosome formation in control and SENP3-knockdown cells using MitoPHfluorin. SENP3 knockdown significantly reduced the levels of SEP fluorescence quenching induced by DFP, indicating a critical role for SENP3 in the disposal of dysfunctional mitochondria (Figure 2B). Taken together, these results suggest that a protein deSUMOylation pathway mediated by SENP3 promotes DFP-induced mitophagy.

**Figure 2.**
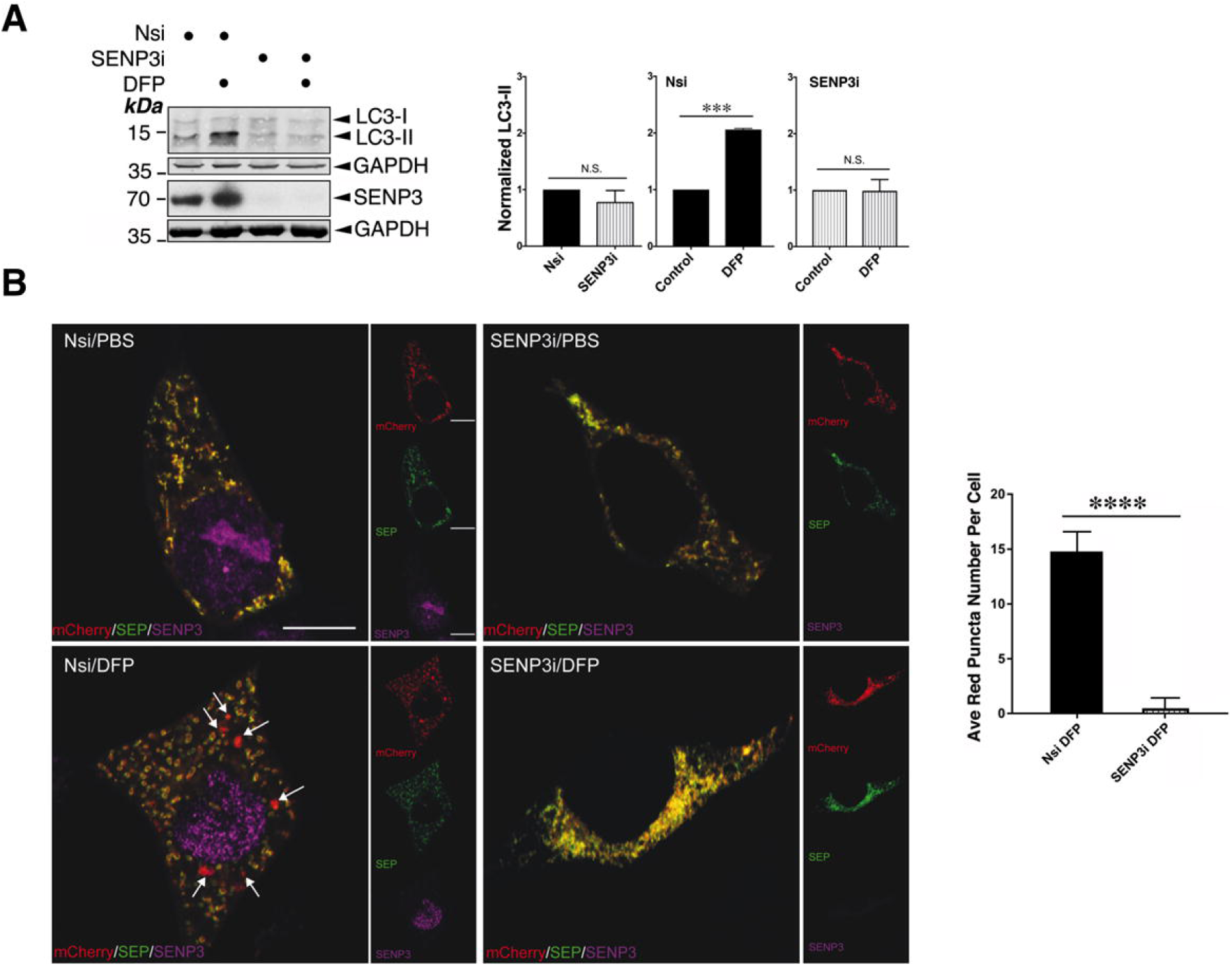
SENP3 is required for mitophagy induced by DFP. A. SENP3 knockdown abolishes DFP-induced LC3-II in HeLa cells. Nsi or SENP3i was transfected into HeLa cells. Two days post transfection and the cells were treated with DFP for a further 24 h. Whole cell lysate samples were blotted as indicated (Nsi, non-specific siRNA; SENP3i, SENP3 siRNA; concentration, 20nM; DFP, 1mM for 24 h). n=3; ***, p<0.001; N.S., not significant, p>0.05; paired t-test. B. SENP3 knockdown abolishes DFP-induced mitophagic autolysosomes. Nsi or SENP3i together with MitoPHfluorin were transfected into HeLa cells in the absence or presence of DFP (24 h), and were analyzed using the MitoPHfluorin construct 72 h post-transfection (Red: mCherry-A or puncta indicating occurrences of mitophagy marked by white arrows; Green: SEP-A; Yellow: mitochondria labelled by mCherry-SEP-A; Magenta: SENP3; Scale bar, 10 μm; n=28 cells per condition; **** p<0.0001; unpaired t-test).

### Fis1 is critical for DFP-induced mitophagy

Since SENP3 regulates mitophagy, we next wondered as to the identity of the specific substrate(s) of its deSUMOylating activity that may mediate this effect. We reasoned that the most likely SENP3 substrates in this pathway are component(s) of the mitochondrial fission machinery. Mitochondrial fission depends on the action of the large GTPase Drp1, a validated SUMO target (Figueroa-Romero, Iniguez-Lluhi et al., 2009, Guo et al., 2013, Prudent, Zunino et al., 2015). Drp1 is recruited to the mitochondrial surface and acts by interacting with specific mitochondrial docking proteins such as Mff, MID49, MID51 and possibly Fis1 in mammalian cells (Loson, Song et al., 2013). However Drp1 does not appear to be essential for mitophagy (Yamashita et al., 2016, Yamashita & Kanki, 2017). Meanwhile, evidence has emerged for the critical role of Fis1 in the mitophagy-mediated disposal of defective mitochondria induced by stressors (Rojansky, Cha et al., 2016, Shen, Yamano et al., 2014, Yamano, Fogel et al., 2014). To investigate if Fis1 is involved in DFP-induced mitochondrial autophagosome formation, we first examined the effect of RNAi-mediated knockdown of Fis1 on the induction of LC3-II in HeLa cells treated with DFP. As expected, Fis1 knockdown eliminates LC3-II induction by DFP in HeLa cells (Figure 3A), indicating a critical role for Fis1 in DFP-induced mitophagic autophagosome formation. We then compared mitochondria-containing autolysosome formation in control and Fis1-knockdown cells using MitoPHfluorin. Fis1 depletion significantly reduced the levels of SEP fluorescence quenching induced by DFP, indicating a critical role for Fis1 in the disposal of dysfunctional mitochondria upon induction of iron-chelation-mediated stress (Figure 3B). Taken together, these results indicate that Fis1 is required for DFP-induced mitophagy.

**Figure 3.**
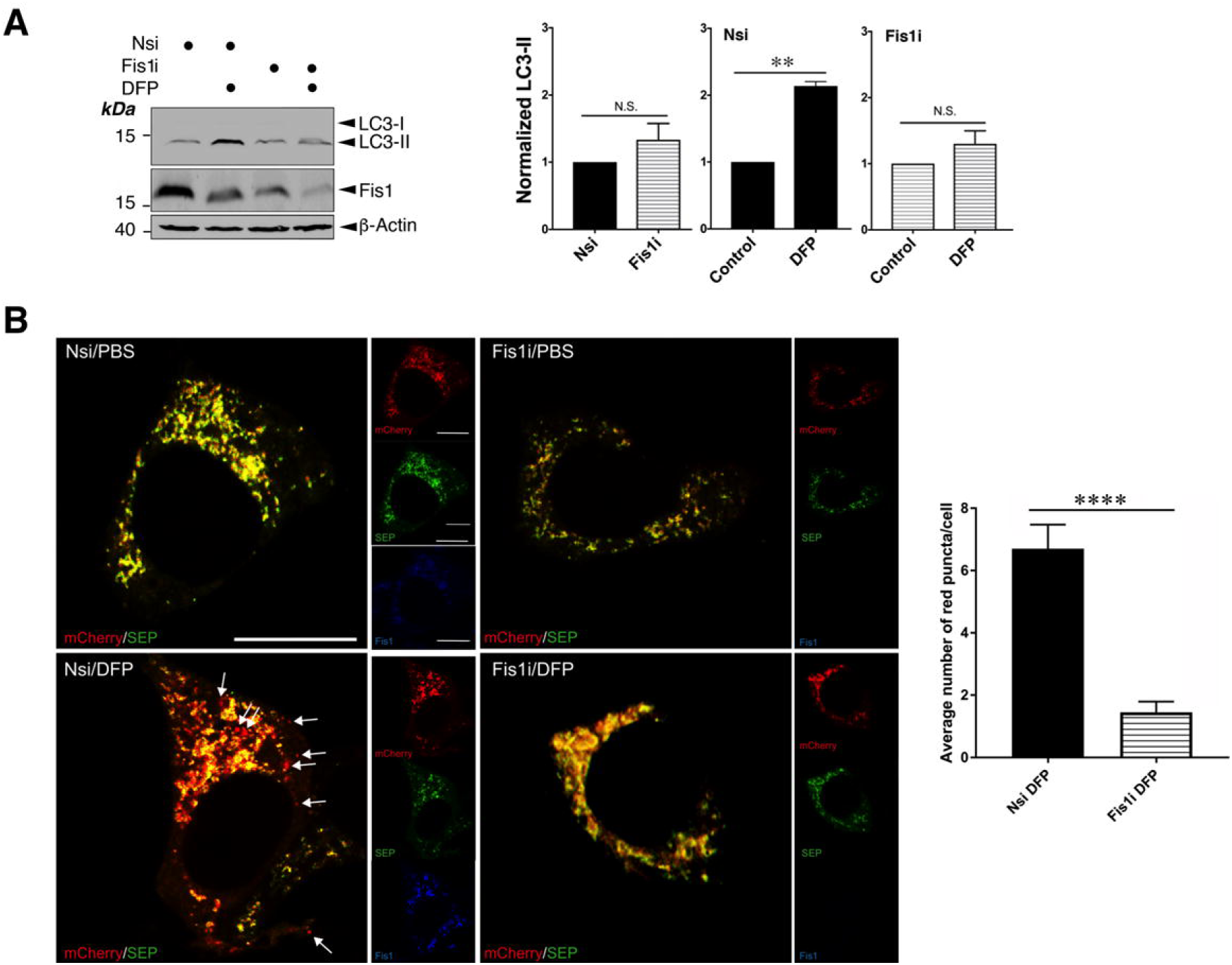
Fis1 is required for DFP-induced mitophagy. A. Fis1 knockdown abolishes DFP-induced LC3-II. Fis1 siRNA was introduced into HeLa cells for 48 h, and the cells were treated with DFP for further 24 h (Fis1i, Fis1 siRNA; concentration, 20nM; DFP, 1mM). Values are presented as mean ± SEM and are normalized to the control value (n=3; **, p<0.01; paired t-test). B. Fis1 knockdown prevents DFP-induced mitophagic autolysosomes. Nsi or Fis1i together with MitoPHfluorin were transfected into HeLa cells in the absence or presence of DFP (24 h), which were analysed using the MitoPHfluorin construct 72 h post-transfection (Red: mCherry-A or puncta indicating occurrences of mitophagy marked by white arrows; Green: SEP-A; Yellow: mitochondria labelled by mCherry-SEP-A; Blue: Fis1; Scale bar, 10 μm; n=20 cells per condition; **** p<0.0001; unpaired-t test).

### Fis1 is a novel SUMO target and is deSUMOylated by SENP3

Since Fis1 had been identified as a potential target for SUMOylation in a proteomics screen for SUMO targets *in vivo* (Tirard, Hsiao et al., 2012), we next verified whether Fis1 was in fact a *bona fide* target for SUMOylation. Flag-Fis1 is efficiently conjugated by His-SUMO-2 in HEK293 cells (Figure 4A). Subsequently under denaturing conditions, we detected conjugation of His-SUMO-2 to endogenous Fis1 in HEK293 cells (Figure 4B). Furthermore, Fis1 antibody-reactive band(s) were detected in immunoprecipitations of endogenous SUMO-2/3 conjugates in HeLa cells (Figure 4C), indicating the existence of SUMOylated endogenously Fis1. Importantly, SENP3 knockdown led to a significant increase in Fis1 SUMO-2-lyation, suggesting SENP3 directly deSUMOylates Fis1 (Figure 4D). Taken together, these findings indicate that Fis1 is subject to SUMO-2/3-lyation under basal conditions, which can be reversed through the action of the deSUMOylating enzyme, SENP3.

**Figure 4.**
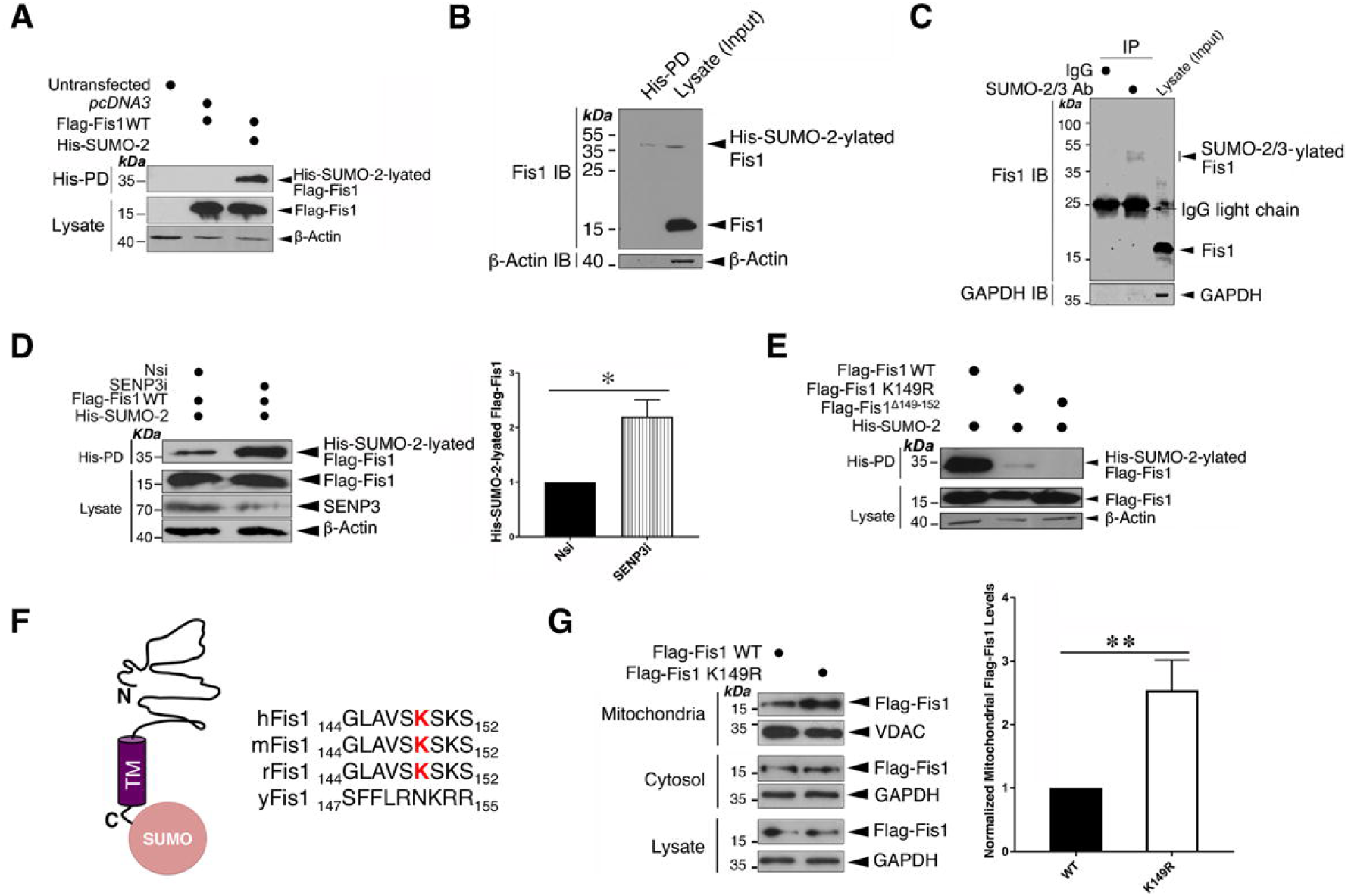
SENP3-mediated deSUMOlyation regulates Fis1 mitochondrial localization. A.Fis1 is a *bona fide* target for SUMOylation. Fis1 is a SUMO target in mammalian cells. Flag-Fis1 together with *pcDNA3* or His-SUMO-2 were transfected into HEK293 cells expressing Ubc9 for 48 h. His-pulldown (His-PD) and lysate samples were detected by immunoblotting for Flag or β-actin. B. Endogenous Fis1 is modified by His-SUMO-2 stably expressed in HEK293 cells. His-pulldown (His-PD) and lysate samples were blotted as indicated. C. Endogenous Fis1 is modified by endogenous SUMO-2/3 in HeLa cells. SUMO-2/3 conjugates were immunoprecipitated (IP) from whole cell lysates prepared from HeLa cells using a SUMO-2/3 antibody. IP and lysate samples were blotted as indicated. D. Fis1 is deSUMOylated by SENP3. SENP3 knockdown enhances Fis1 SUMOylation. Nsi or SENP3i were transfected into HEK293 cells expressing Flag-Fis1, His-SUMO-2 and Ubc9 for 48 h. His-pulldown and lysate samples were detected by immunoblotting for Flag, SENP3 or β-actin (n=3; * p<0.05; paired-t test). E. Amino acids including lysine 149 at the C-terminus of Fis1 are required for Fis1 SUMOylation. Flag-Fis1, Flag-Fis1 K149R mutant or Flag-Fis1 Δ149-152 mutant together with His-SUMO-2 were transfected into HEK293 cells expressing Ubc9 for 48 hours. His-pulldown and lysate samples were blotted as indicated. F. Alignment (the right panel) shows amino acid sequences of the C-terminal tail of Fis1 from the indicated organisms: human (h), mouse (m), rat (r) and *S. cerevisiae* (y); red letters mark the lysine residue required for Fis1 SUMOylation. Schematic (the left panel) illustrates of SUMO conjugation at the C-terminal tail of Fis1 (N, N-terminus; C, C-terminus). G. The SUMOylation status of Fis1 influences its mitochondrial localization. Flag-Fis1 WT or Flag-Fis1 SUMOylation deficient mutant K149R was expressed in HeLa cells. Whole cell lysates, and cytosolic and mitochondrial fractions were prepared and blotted as indicated (the upper panel). Histograms (the right panel) show normalized levels of Flag-Fis1 associated with the mitochondrial fraction (Value are presented as mean ± SEM and are normalized to the control value; n=7; **, p<0.01; paired t-test).

### Fis1 is primarily SUMOylated at lysine 149

To identify which lysine in Fis1 is SUMOylated, we performed site-directed mutagenesis. Fis1 contains one high-probability consensus lysine residue (K119) and three high-probability non-consensus lysine resides (K67, K149 and K151), predicted by SUMOsp (Xue, Zhou et al., 2006). While mutation of either the K119 or the K67 to a non-SUMOylatable arginine (R) did not reduce Fis1 SUMOylation in a HEK293 cell-based assay under denaturing conditions (Supplementary Figure 2A), K149R mutation in Fis1 reduces Fis1 SUMOylation by ∼90% (Figure 4E). Moreover, the deletion of the final four amino acid residues (^149^**K**S**K**S^152^) completely eliminated Fis1 SUMOylation (Figure 4E). This finding led us to question if K151 in Fis1 was a minor site for SUMO conjugation. Unexpectedly, K151R mutation does not seem to reduce Fis1 SUMOylation, however it is possible that this site becomes a target only in the absence of the nearby K149 (Supplementary Figure 2B). These findings indicate that K149 is the major SUMO acceptor site in Fis1 (Figures 4F).

### The SUMOylation status of Fis1 acts as a key switch to regulate its mitochondrial localization and mitophagy induction

Interestingly, K149 is known to be critical for localizing Fis1 to mitochondria. Non-conservative mutation of K149 to alanine (A) greatly reduces Fis1 mitochondrial localization (Delille & Schrader, 2008, Stojanovski, Koutsopoulos et al., 2004) whilst Fis1 with the conservative K149R mutation is localized to mitochondria (Alirol, James et al., 2006). However, whether the K149R mutation results in any changes in the levels of mitochondrial Fis1 remained undetermined. According to our quantitative data, the K149R mutation in fact increases the levels of Fis1 in mitochondria (Figures 4G), suggesting that deSUMOylation promotes Fis1 mitochondrial localization. We then asked if changes in Fis1 SUMOylation have an impact on mitophagy. We examined the effect of expressing Fis1 wild-type (WT) or Fis1 K149R mutant in HeLa cells, where endogenous (human) Fis1 was depleted by siRNA (Supplementary Figure 3), on LC3-II induction. Fis1 mutant-expressing cells show elevated levels of LC3-II, in the absence of DFP (Figure 5A). Taken together, these findings suggest that the SUMOylation status of Fis1 serves as a key molecular switch to regulate its mitochondrial localization and mitophagic autophagosome formation.

**Figure 5.**
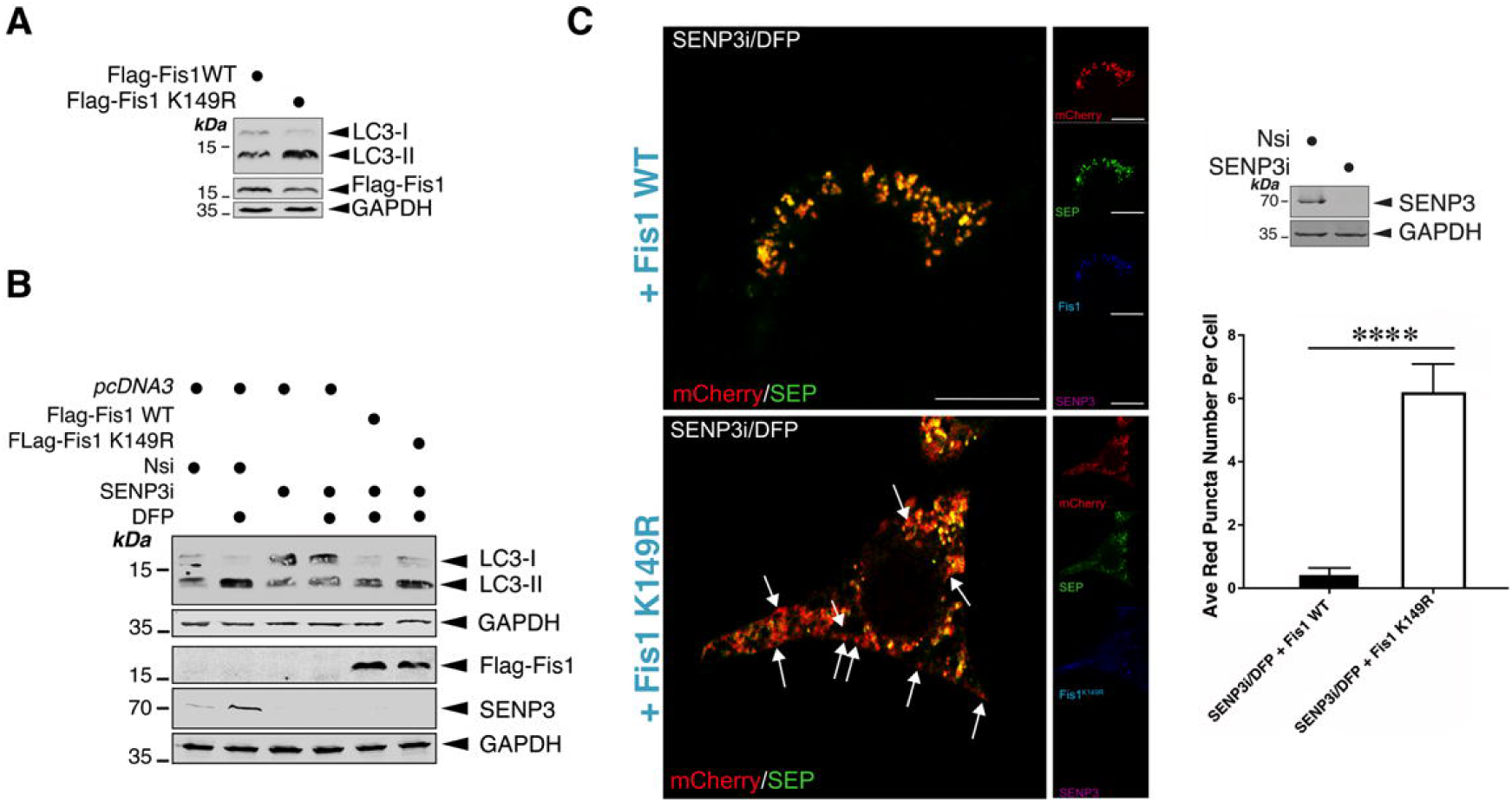
SUMOylatable Fis1 is essential for SENP3 regulation of mitophagy. A. Expression of a SUMOylation-deficient Flag-Fis1 K149R mutant in HeLa cells induces LC3-II. Flag-Fis1 WT or Flag-Fis1 K149R mutant were transfected into HeLa cells in which endogenous Fis1 was depleted using siRNA for 48 h. Lysate samples were blotted as indicated. B. Expressing a SUMOylation-deficient Fis1 mutant reverses the effect of SENP3 knockdown on DFP-induced LC3-II induction. *pcDNA3*, Flag-Fis1 WT or Flag-Fis1 K149R mutant were transfected into HeLa cells in which SENP3 was depleted using siRNA for 48 h, and the cells were treated with DFP for a further 24 h. Lysate samples were blotted as indicated. C. SENP3 depletion does not abolish mitophagic puncta detected in HeLa cells expressing SUMOylation deficient CFP-Fis1 K149R in the presence of DFP. CFP-Fis1 WT or CFP-Fis1 K149R mutant were transfected into HeLa cells in which SENP3 was depleted using siRNA for 48 h, and the cells were treated with DFP for a further 24 h (Red: mCherry-A or puncta indicating occurrences of mitophagy marked by white arrows; Green: SEP-A; Cyan/blue: CFP-Fis1; Scale bar, 10 μm; n=12 cells per condition; **** p<0.0001; unpaired t-test). Knockdown of SENP3 was confirmed by immunoblotting (the upper right panel).

### SENP3 deSUMOylates Fis1 to regulate DFP-induced mitophagy

The obove-mentioned findings led us to hypothesize that SENP3-mediated deSUMO-2/3-ylation of Fis1 would be a critical step DFP-induced mitophagy. To test this hypothesis, we examined the levels of LC3-II in DFP-treated HeLa cells in which SENP3 was knocked down and Flag-Fis1 WT or Flag-Fis1 K149R mutant was expressed. As expected, upon DFP treatment, higher levels of LC3-II were associated with SENP3-knockdown cells expressing the SUMOylation-deficient K149R mutant (Figure 5B). Since SENP3 knockdown blocks LC3-II expression only in the presence of SUMOylatable FIs1, these results indicate that Fis1 is the downstream deSUMOylation target for SENP3-mediated mitophagic autophagosome formation. In parallel, we monitored levels of SEP quenching of the MitoPHfluorin reporter in HeLa cells in which SENP3 was knocked down and CFP-Fis1 WT or CFP-Fis1 K149R mutant was expressed. As predicted, knockdown of SENP3 did not reduce mitophagic autolysosome formation in cells expressing Fis1 K149R mutant in the presence of DFP (Figure 5C), indicating the importance of the SUMOylation status of Fis1 for SENP3-mediated mitophagic autolysosome formation. Taken together, these results have verified our hypothesis that SENP3-mediated deSUMO-2/3-ylation of Fis1 is required for this previously uncharacterized mitophagy pathway.

## Discussion

Healthy mitochondria are vital to generate energy for eukaryotic cells and accumulation of dysfunctional mitochondria has been linked to ageing and age-related diseases. This age-related pathological feature is believed to be due to defective mitophagy, which mediates mitochondrial quality control via lysosomal degradation. However, the mechanisms that regulate mitophagy and the key pathways involved in mitochondrial quality control remain largely unknown. Here we report a previously uncharacterized pathway that regulates mitophagy via modulation of the SUMOylation status of the mitochondrial protein Fis1 by the SUMO-2/3-specific protease SENP3 (Figure 6). To the best of our knowledge, this is the first example showing functional crosstalk between protein SUMOylation and regulation of mitophagy.

**Figure 6.**
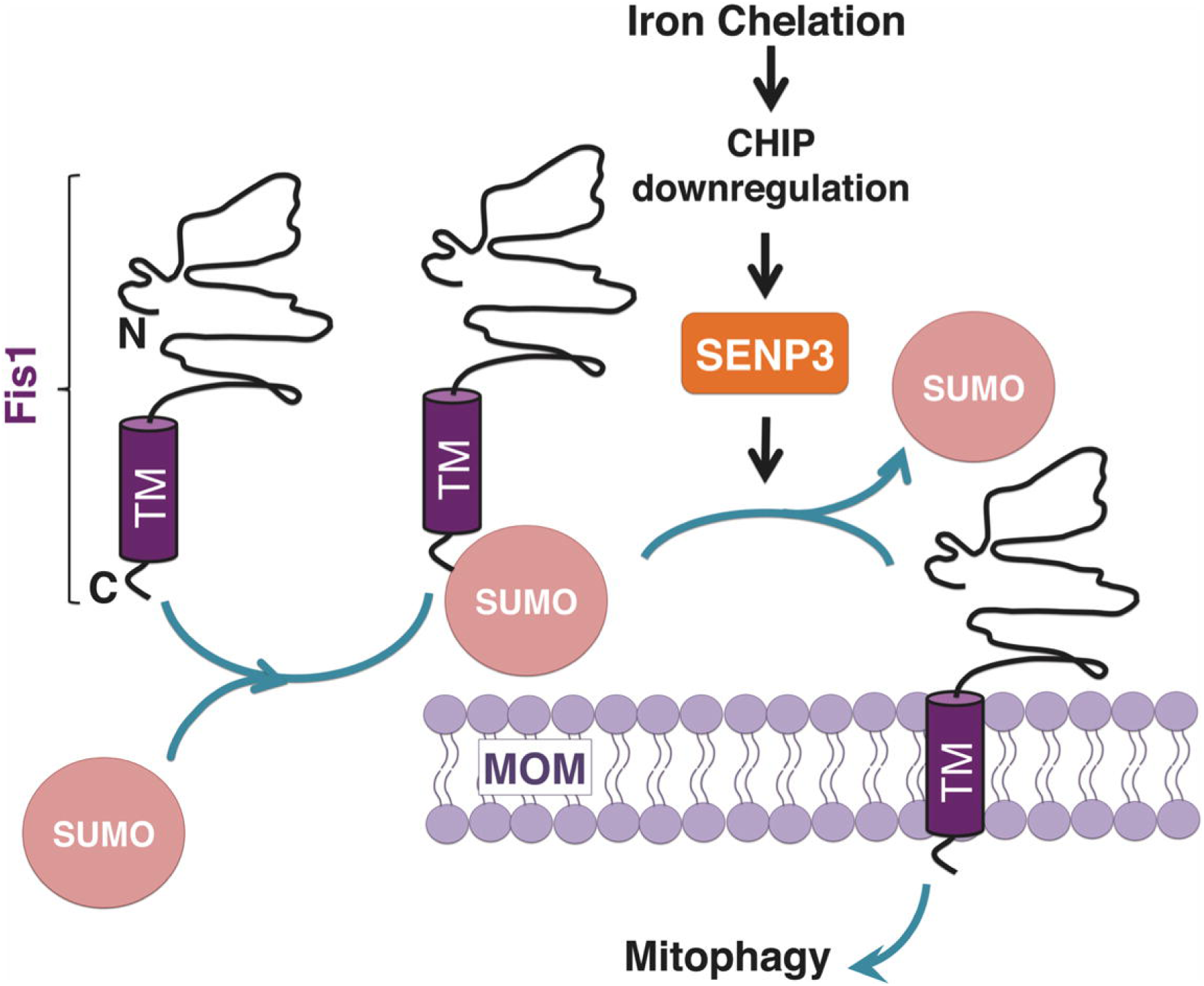
Schematic shows the roles of SUMOylation and SENP3-mediated deSUMOylation in Fis1 mitochondrial localization. DFP treatment enhances SENP3-mediated deSUMOylation of Fis1, thereby facilitating Fis1 mitochondrial localisation that is required for mitophagy.

### Modulation of SENP3 stability by mitophagy induction

In mammals the Ub-proteasome system (UPS) controls SENP3 levels through proteasome-mediated degradation (Kuo, den Besten et al., 2008, Yan et al., 2010). It has been shown previously that the E3 Ub ligase CHIP mediates UPS-dependent degradation of SENP3 (Yan et al., 2010). In this present study, we found that upon DFP-mediated iron chelation SENP3 is stabilised whilst CHIP levels are reduced, suggesting that CHIP reduction may lead to SENP3 stabilisation through a reduction in SENP3 degradation.

A key finding of this study is that SENP3 is pivotal for mitophagy induction by iron chelation. SENP3 stabilisation is temporally correlated with decreased levels of global SUMO-2/3 conjugation. Consistent with this, knockdown of SENP3 abolishes DFP-induced formation of both mitophagic autophagosomes and autolysosomes, further suggesting that sufficient levels of SENP3 are required for the occurrence of mitophagy upon iron chelation. We further identify Fis1 as a key target for SENP3-mediated deSUMOylation in mitophagy regulation since expression of a SUMOylation-deficient mutant of Fis1 abolishes the effect of SENP3 knockdown on DFP-induced mitophagy.

### SUMOylation and deSUMOylation of Fis1 and Fis1 mitochondrial localization

The canonical mitophagy pathway involves Pink1 and Parkin. However, mice lacking either Pink1 or Parkin show very mild defects in mitochondrial function (Whitworth & Pallanck, 2017). In contrast, genetic depletion of Fis1 in mammalian cells blocks mitophagy-mediated disposal of dysfunctional mitochondria (Rojansky et al., 2016, Shen et al., 2014, Yamano et al., 2014), signifying a major role for Fis1 in mitochondrial quality control. However, it remains unclear how Fis1 mediates mitophagy. Emerging evidence indicates that mitophagy induction requires Fis1 mitochondrial localization (Rojansky et al., 2016). In keeping with the importance of Fis1 mitochondrial localization, our results showing that mutation of the major SUMOylated lysine (K149) enhances its mitochondrial presence indicate that preventing SUMOylation promotes Fis1 mitochondrial localization and suggest that SUMO-2/3 modification may act as a molecular switch to regulate levels of mitochondrial Fis1. However, it should be noted that, in fact, at any given time only a small proportion of Fis1 is SUMOylated. This argues against constitutive SUMOylation being required to maintain Fis1 in the cytosol. Rather, similar to what we have previously suggested about the roles for SUMOylation in Drp1 mitochondrial partitioning (Guo et al., 2013), deSUMOylation may act as a ‘mobilisation factor’ for Fis1 mitochondrial targeting.

In conclusion, we define a previously unidentified SUMO-dependent pathway in which UPS-dependent stabilisation of SENP3 upon iron chelation enhances mitochondrial targeting of Fis1 to facilitate mitophagy induction. This pathway represents a novel example of a target-specific function of SENP3 in a mitochondrial quality control programme and reveals a potential therapeutic target for restoring impaired or defective mitophagy in health and disease.

## Materials and methods

### Cloning and mutagenesis

DNA constructs encoding Flag-SENP3, His-SUMO-2, and GST-(mouse) Fis1 were described previously (Guo et al., 2013, Guo et al., 2017). Flag-Fis1 was generated by insertion of the relevant cDNA amplified from GST-Fis1 into the BamH1/Not1 sites of pcDNA3. Flag-Fis1 mutants, K67R, K119R, K149R, K151R and Δ149-152, were made by PCR-based mutagenesis. CFP-Fis1 was generated by sequential insertions of cDNA encoding CFP and cDNA encoding mouse Fis1 into the BamHI/EcoRI sites and EcoRI/NotI sites of pcDNA3, respectively. To visualise ‘mitophagic flux’, a novel probe (MitoPHfluorin) was generated as follows: cDNA sequences encoding Super-ecliptic pHluorin (SEP, a pH sensitive GFP variant) and the mitochondrial target sequence (LILAMLAIGVFSLGAFIKIIQLRKNN, termed as ‘A’) of the ActA protein from *Listeria monocytogenes* were amplified using PCR from SEP-TOPO (Ashby et al., 2004) and GFP-A (Guo et al., 2017), respectively. The plasmid encoding the fusion protein mCherry-SEP-A was then made by sequential insertion of the cDNA encoding SEP and the cDNA encoding A into a pmCherry-C3 construct.

### Cell culture

HeLa and HEK293 cells were cultured in Dulbecco’s modified Eagle’s medium (DMEM; Lonza) containing 10% fetal bovine serum (FBS), 5 mM glutamine, and 100 units/ml penicillin/streptomycin at 37°C in humidified air supplemented with 5% CO_2_, as previously described (Guo et al., 2013, Guo et al., 2017). HEK293 N3S cells stably expressing 6His-SUMO-2^T90K^ (Tammsalu, Matic et al., 2014) were a gift from Hay and were cultured in DMEM containing 10% FBS, 5□mM glutamine, 100□units/ml penicillin/streptomycin in the presence of 1 μg /ml puromycin.

### DFP treatment for mitophagy induction

Confluent HeLa cells grown in 6- or 12-well plates were treated with deferiprone (DFP; 379409, Sigma-Aldrich; working concentration, 1mM) for the time durations as indicated.

### DNA and siRNA transfections

For DNA, siRNA, or DNA-siRNA co-transfection into HeLa or HEK293 cells, we used jetPRIME reagents (Polyplus Transfection) as previously described, according to the manufacturer’s instructions, and used the cells within 72 h. siRNA duplexes used were as follows: human SENP3 siRNA (sc-44451, Santa Cruz), non-specific siRNA or a pool of human Fis1 siRNA (synthesized by Eurofins MWG Operon). The sequences chosen to silence Fis1 are CUACCGGCUCAAGGAAUAC and GGAAUACGAGAAGGCCUUA. The two sequences are not present in mouse Fis1, therefore, plasmids encoding mouse Fis1 are naturally resistant to the siRNA targeted against human Fis1.

### Subcellular fractionation

Cytosol and mitochondria fractions were prepared from HeLa cells using a Cell Fractionation Kit (Abcam), according to the manufacturer’s instructions.

### Preparation of cell lysate samples and immunoprecipitation

Treated/transfected HeLa or HEK293 cells were washed once with ice-cold phosphate-buffered saline and then lysed in a buffer containing 20 mM Tris, pH 7.4, 137 mM NaCl, 25 mM β-glycerophosphate, 2 mM sodium pyrophosphate, 2 mM EDTA, 1% Triton X-100, 10% glycerol, and 1x cOmplete™ Protease Inhibitor Cocktail (Roche). N-Ethylmaleimide (NEM; 20mM) was added when protein samples were prepared to assess effects of SUMOylation/deSUMOylation. Following sonication, insoluble material was removed from lysates by centrifugation for 15 min at 16,000 × g at 4°C. In experiments to detect endogenous Fis1 SUMO-2/3-lyation, cells were lysed in the above-mentioned buffer plus 1% SDS, and heated at 95°C for 5□min. The lysate samples were then diluted 1: 10 in the same buffer without SDS in the presence of 20□mM NEM. Using SUMO-2/3 antibody (Clone 1E7, MBL) endogenous SUMO-2/3 conjugates from the lysate samples were immunoprecipated as described previously (Guo et al., 2017).

### Histidine pull-down

His-SUMO-2 conjugates were purified through histidine pulldowns (His-PD) under denaturing conditions as previously described (Guo et al., 2013, Guo et al., 2017). Briefly, transfected HEK293 cells or HEK293 N3S cells stably expressing 6His-SUMO-2^T90K^, were immediately lysed in a denaturing buffer containing 6 M guanidinium-HCl, 0.1 M Na_2_HPO_4_/NaH_2_PO_4_, 0.01 M Tris-HCl, pH 8.0, sonicated, and incubated with Ni^2+^–NTA beads (Qiagen) for 3 h at room temperature. Following the incubation, the nickel beads were washed twice with the denaturing buffer, two times with wash buffer containing 8 M urea, 0.1 M Na_2_PO_4_/NaH_2_PO_4_, 0.01 M Tris-HCl, pH 6.3, and twice with PBS. Bound proteins were eluted in 6× SDS loading buffer and resolved by SDS–PAGE. His-SUMO-2 conjugates were immunoblotted using an antibody against Flag or Fis1.

### Immunoblotting

Lysate, IP or His-PD Samples were resolved by SDS-PAGE (10-15% gels) and transferred to Immobilon-P membranes (Millipore Inc.), which were then immunoblotted with the following antibodies against: β-actin (Sigma), CHIP (Cell Signaling), Fis1 (Proteintech), Flag (Proteintech), GAPDH (Santa Cruz biotechnology), GST (GE Healthcare), LC3-II (Cell Signaling), SENP3 (Cell Signaling), SENP5 (Abcam). SUMO-1 (Santa Cruz biotechnology), SUMO-2/3 (Cell Signaling; 1E7, MBL) or Tom20 (Santa Cruz biotechnology). Immune complexes were detected using either HRP-conjugated secondary antibodies (Sigma) followed by enhanced chemiluminescence (GE Healthcare) or using fluorescent secondary antibodies (LI-COR). To avoid the potential interference from mouse IgG heavy chain, a HRP-conjugated VeriBlot secondary antibody (ab131366, Abcam) was used in immunoblotting for SUMO-2/3-lysated endogenous Fis1 in IP samples.

Each immunoblot presented is representative of at least three experiments carried out using different cell populations.

### Fluorescence and microscopy imaging

HeLa cells were plated on 35mm glass bottom dishes and transfected after 24 hr. After 48 hr, cells were treated with DFP (1 mM) or PBS for a further 24 hr. Cells were fixed for 12 min at RT in 4% paraformaldehyde/PBS. Following three 5 min-washes with PBS to remove residual PFA, cells were permeabilised in 0.3% Triton X-100/PBS for 15 min and then blocked for 1 hr in 0.01% Triton X-100, 0.3% Fish skin gelatin/PBS. Antibodies were diluted in blocking buffer and incubated for 1.5 hr at RT. Immunostaining was performed using the following antibodies: SENP3 (Cell signalling, 1:400), Fis1 (Proteintech, 1:400), Flag (Proteintech, 1:500). Cells were analysed with either an inverted fluorescence microscope (Axiovert 200 M; Carl Zeiss Microimaging, Inc.) or a confocal microscope (Zeiss LSM 880 Airyscan) within the Wolfson Light Microscope Facility at the University of Sheffield.

### Statistics

For comparison of conjugates of SUMO-2/3 or SUMO-1, and levels of SENP3, SENP5, CHIP, or LC3-II in lysate or His-PD samples between time-matched two groups, paired Student’s test with two-tail p-value was performed. For analysis of mitophagic autolysosomes, unpaired Student’s t-tests with two-tail p-value was performed comparing mean of red punctum numbers. Following quantification using ImageJ software, levels of proteins of interest were normalized to β-actin or GAPDH control levels. Each value is presented as mean ± SEM and is expressed as percentage of control value. To avoid bias all experiments were performed blind.

## Acknowledgments

Start-up fund (BMS314826 to CG), Royal Society Research Grant (157589-11-1 to CG) and a PhD studentship (BMS 315637 to EW) from the Department of Biomedical Science at the University of Sheffield supported this work. We are also grateful to BBSRC (BB/R00787X) and Parkinson’s UK for financial support (to KAW and REC, respectively). We thank C Smythe for critical reading of the manuscript and great logistical support, and EH Hettema for generous technical help with initial imaging experiments. We thank JM Henley for supplying plasmids and RT Hay for supplying the HEK293 N3S cells.

## Author contribution

CG and EW conceived the project with valuable input from KAW. EW and CG designed and performed all biochemical and molecular biological analysis, EW performed all cell imaging assays in cell culture, and KAW and REC conducted some cell culture and sample preparation work. CG provided project management and wrote the manuscript with hypothesis development, experimental design and data interpretation contributed by all authors. The funders had no role in study design, data collection and analysis, decision to publish, or preparation of the manuscript.

## Supplementary figure legends

**Supplementary figure 1.**
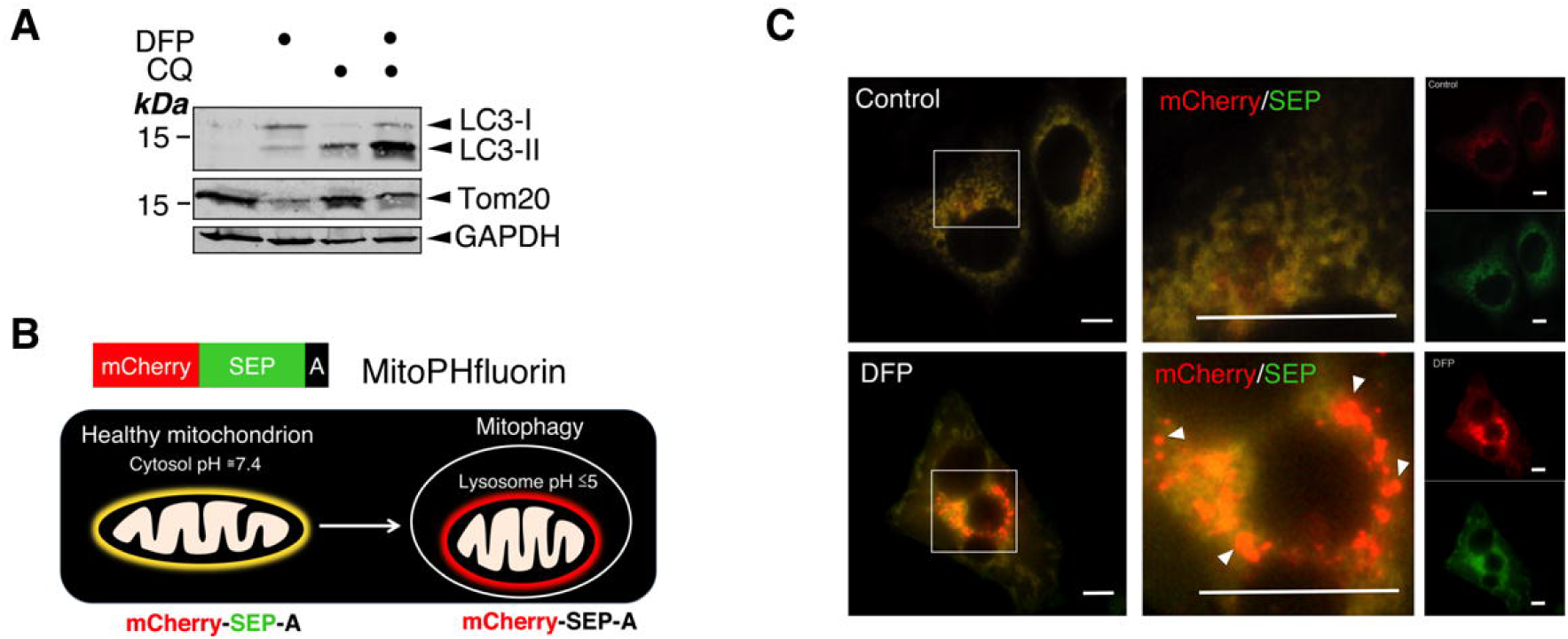
Characterization of iron chelation-induced mitophagy. A. DFP (1mM for 24 h) induces LC3-II in HeLa cells. As a widely used autophagy inhibitor, chloroquine (CQ; 50 μM for 12 h) was used to increase the visualization of the autophagosome marker LC3-II. B. Schematic illustrates i) MitoPHfluorin, a tandem-tagged construct encoding mCherry-SEP-A: SEP, Super-ecliptic pHluorin (a pH-dependent GFP variant) and the mitochondrial targeting sequence derived from the ActA protein fused to the C-terminus of SEP, and ii) a mitophagy assay using the MitoPHfluorin construct. Under basal conditions, in green/red merged images mitochondria are labelled as yellow structures whilst upon mitophagy, mitochondria are sequestered within autolysosomes where fluorescence of SEP is quenched due to the acidic environment and therefore outlined as red structures. C. Detection of DFP-induced mitophagic autolysosomes using MitoPHfluorin. MitoPHfluorin was transfected into HeLa cells in the absence or presence of DFP (1mM for 24 h), and cells analysed 48 h post-transfection (Scale bar is 10 μm; occurrences of mitophagy marked by arrow heads).

**Supplementary figure 2.**
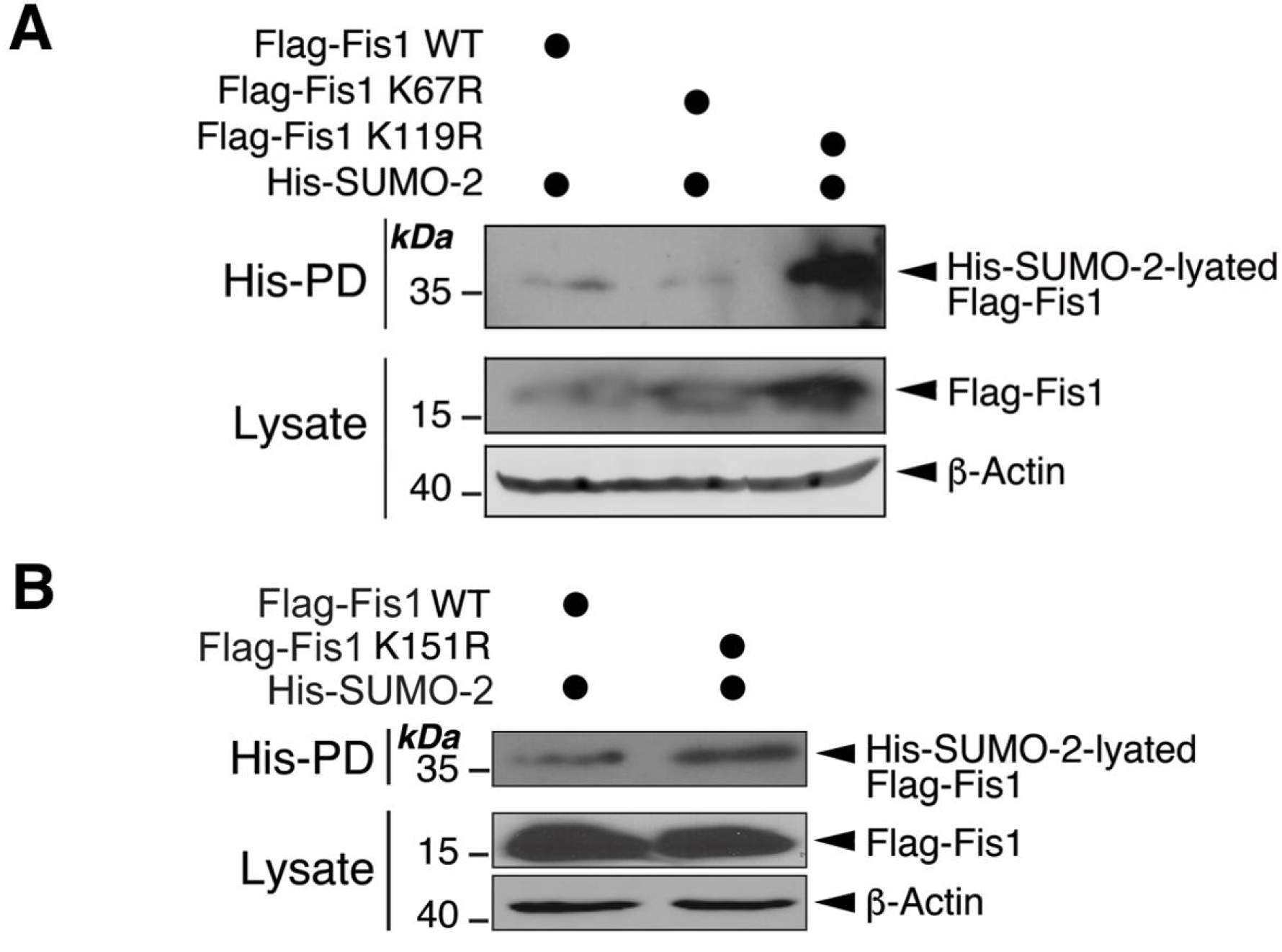
Mutation of lysine 67, lysine 119 or lysine 151 to arginine does not reduce Fis1 SUMOylation. A. Flag-Fis1 WT, Flag-Fis1 K67R or Flag-Fis1 K119R mutant together with His-SUMO-2 were transfected into HEK293 cells expressing Ubc9 for 48 h. B. Flag-Fis1 WT, or Flag-Fis1 K151R mutant together with His-SUMO-2 were transfected into HEK293 cells expressing Ubc9 for 48 h. In (A) and (B), His-pulldown and lysate samples were detected by immunoblotting for Flag or β-actin.

**Supplementary figure 3.**
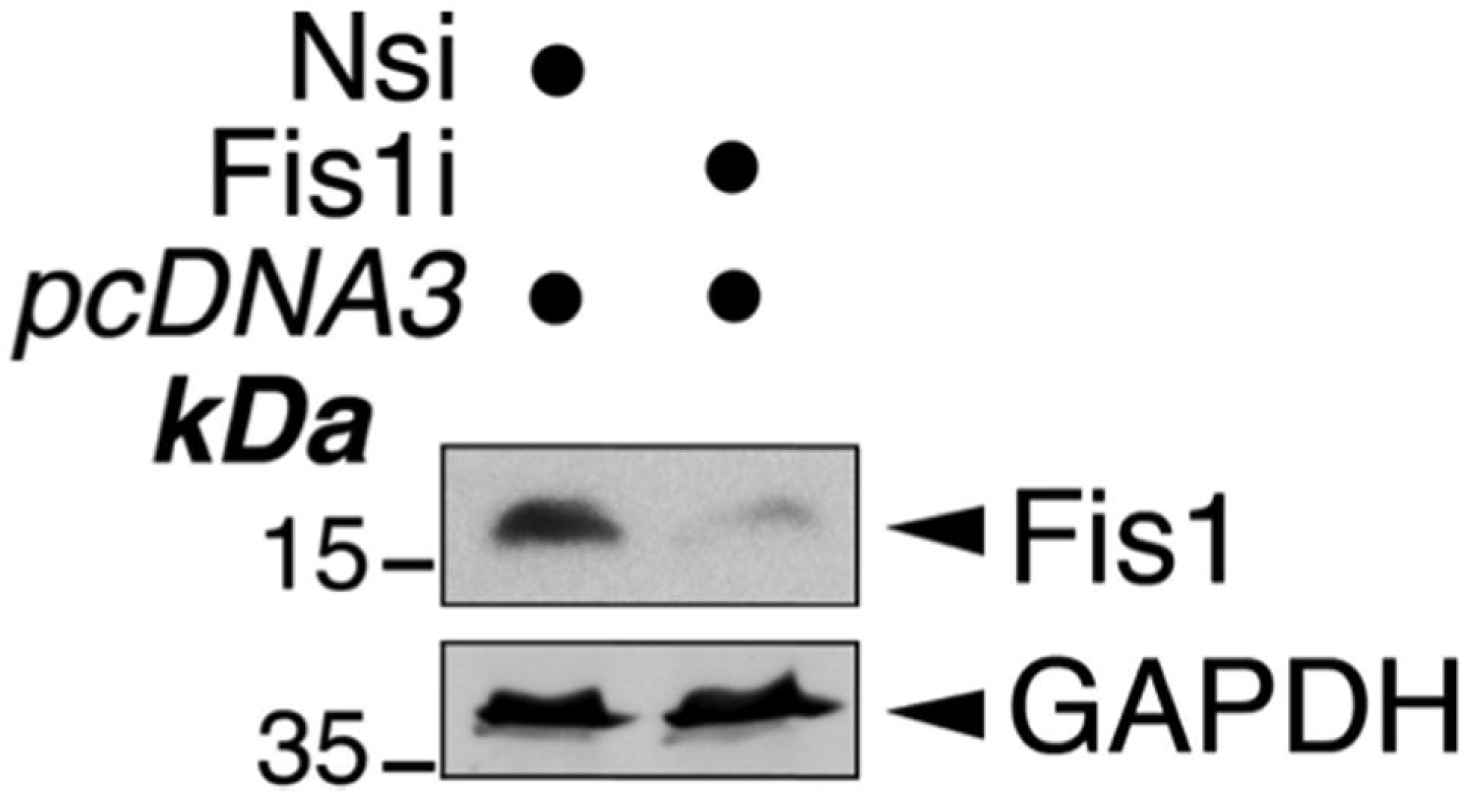
Knockdown of Fis1 in HeLa cells. *pcDNA3* control (for Flag-Fis1 or Flag-Fis1 K149R) was transfected into Fis1 knockdown HeLa cells for 48 h. Knockdown of Fis1 was confirmed by immunoblotting.

